# Cofactors revisited – predicting the impact of flavoprotein-related diseases on a genome scale

**DOI:** 10.1101/343624

**Authors:** Agnieszka B. Wegrzyn, Sarah Stolle, Rienk A. Rienksma, Vítor A. P. Martins dos Santos, Barbara M. Bakker, Maria Suarez-Diez

**Affiliations:** Systems Medicine of Metabolism and Signaling, Laboratory of Pediatrics, University Medical Center Groningen, University of Groningen, 9713 AV Groningen, The Netherlands; Systems Biology Centre for Energy Metabolism and Ageing, University of Groningen, 9713 AV Groningen, The Netherlands; Systems and Synthetic Biology, Wageningen University & Research, 6708 WE Wageningen, The Netherlands; Lifeglimmer GmbH., 12163 Berlin, Germany

**Keywords:** FAD, FMN, flavoprotein, inborn errors of metabolism, human genome-scale reconstruction, constraint-based modelling

## Abstract

Flavin adenine dinucleotide (FAD) and its precursor flavin mononucleotide (FMN) are redox cofactors that are required for the activity of more than hundred human enzymes. Mutations in the genes encoding these proteins cause severe phenotypes, including a lack of energy supply and accumulation of toxic intermediates. Ideally, patients should be diagnosed before they show symptoms so that treatment and/or preventive care can start immediately. This can be achieved by standardized newborn screening tests. However, many of the flavin-related diseases lack appropriate biomarker profiles. Genome-scale metabolic models can aid in biomarker research by predicting altered profiles of potential biomarkers. Unfortunately, current models, including the most recent human metabolic reconstructions Recon and HMR, typically treat enzyme-bound flavins incorrectly as free metabolites. This in turn leads to artificial degrees of freedom in pathways that are strictly coupled. Here, we present a reconstruction of human metabolism with a curated and extended flavoproteome. To illustrate the functional consequences, we show that simulations with the curated model – unlike simulations with earlier Recon versions - correctly predict the metabolic impact of multiple-acyl-CoA-dehydrogenase deficiency as well as of systemic flavin-depletion. Moreover, simulations with the new model allowed us to identify a larger number of biomarkers in flavoproteome-related diseases, without loss of accuracy. We conclude that adequate inclusion of cofactors in constraint-based modelling contributes to higher precision in computational predictions.

## 1. Introduction

In the past decade, systems biology modelling has become indispensable to explore the behaviour of metabolic networks and gain insight into their response to disease mutations. Genome-scale models of metabolism comprise the entire set of biochemical reactions known to exist in an organism or cell type of interest (as described in depth in [1]). These models represent the metabolic potential of living systems and provide a comprehensive framework to understand metabolism. Genome-scale models integrate biochemical and genotypic data and enable efficient exploration of associations between genotypes and phenotypes, and mechanisms of (patho)physiology [2]. The most recent consensus and generic human metabolic reconstructions are Recon 2.2 [3], Recon 3D [4], and HMR 2.0 [5]. Both are comprehensive models aiming to describe all known metabolic reactions within the human body.

Genome-scale constraint-based models contain mass-balanced chemical equations for each metabolic reaction in a specific cell type or organism [6]. Classically, neither enzyme kinetics nor enzyme concentration are accounted for in this approach [7]. The recently published GECKO method [8] for yeast models, which links enzyme abundances with reaction fluxes addresses this issue. Similarly, Shlomi et al. [9] successfully studied the Warburg effect in human genome-scale model by taking an enzyme solvent capacity into consideration. However, general incorporation of enzyme concentrations and kinetics in human models remains to be addressed. Consequently, taking enzyme-bound cofactors into account remains a challenge.

Cofactors are molecular compounds required for the enzyme’s biological activity. Chemically, they can be divided into two groups: inorganic ions that are taken up by the cell from the environment, and more complex organic or metalloorganic molecules – also called coenzymes - which are (partly) synthesized in the cell. The latter are often derived from vitamins and organic nutrients and their biosynthesis pathways must be covered by genome-scale models.

The flavins FAD and FMN are redox cofactors required for the activity of 111 human enzymes, 52 of which are known to cause human diseases if inactive [10]. In human cells, flavins are synthesized from their precursor riboflavine, also known as vitamin B2. Flavins exist in three different redox states, namely a quinone, semiquinone and hydroquinone state. Unlike nicotinamide adenine dinucleotide (NAD) which diffuses freely between enzymes, flavins are bound to enzymes [11]. Enzymes that require bound FAD or FMN for their enzyme activity are called flavoproteins.

A classical flavoprotein-dependent disease is multiple-acyl-CoA-dehydrogenase deficiency (MADD), also known as glutaric aciduria type II. It is caused by a defect in one of the electron transfer flavoproteins (encoded by *ETFA* or *ETFB* genes) or in the ‘electron transfer flavoprotein ubiquinone oxidoreductase’ (ETF-QO, encoded by *ETFDH* gene) [12]. These FAD-containing enzymes are crucial to link both mitochondrial fatty acid oxidation (mFAO) and amino acid metabolism (mostly that of sarcosine and dimethylglycine) to the mitochondrial respiratory chain. Depending on the residual ETF or ETF-QO activity, MADD may lead to a life-threatening lack of energy supply to the body, with episodes of severe metabolic decompensation, hypoglycaemia, metabolic acidosis, sarcosinemia and cardiovascular failure. The available treatment consists of low-fat, low-protein and high-carbohydrate diet with riboflavin, glycine and L-carnitine supplementation. However, this treatment is not effective for neonatal patients [13] for whom experimental treatment with sodium-D,L-3-hydroxybutyrate showed promising results [14]. To screen for and diagnose diseases, as well as provide a prognosis for disease severity and treatment outcome, biomarkers are used. According to the NIH definition, a biomarker is “a biological molecule found in blood, other body fluids, or tissues that is a sign of a normal or abnormal process, or of a condition or disease” [15]. They are measured routinely in a non-invasive manner in plasma, urine or faeces [16]. Several inborn errors of metabolism (IEM) have their metabolic phenotypes characterized [17]. However, for many diseases no known biomarker profile exists [18] or their sensitivity is low causing some patients to be wrongly diagnosed in the screening process [19]. Additionally, clinical presentations of IEM are often non-specific and are described as a spectrum rather than a clear one gene – one phenotype – one disease relation. This further justifies the search for more sensitive and accurate biomarkers. Systems biology approaches have already been proven to aid in such research by revealing known and novel biomarkers of several IEMs [17, 20–23]. Human genome-scale models, Recon 2 and HMR 2, can be used to identify biomarkers from altered patterns of metabolites taken up or excreted by tissues [24]. These models typically treat flavins and other enzyme-bound cofactors as free-floating metabolites, as seen on a Fig. 1A and B [5,23]. This allows enzymes to inadvertently oxidize or reduce flavin molecules that are in reality confined to another enzyme. In the models, this leads to artificial uncoupling of pathways. Such an artefact is seen clearly in the case of MADD where electrons from mFAO should be coupled to the oxidative phosphorylation system by the ETF/ETF-QO proteins (Fig. 1C). In the current models, however, FADH produced from mFAO can also be re-oxidized via other pathways: by reversal of the mitochondrial succinate dehydrogenase reaction as well as an artificial reaction that represents triglycerides synthesis (Fig. 1B, model Recon2.2). Another artefact in Recon2.2, but not HMR 2, is that flavin biosynthesis is not explicitly required due to the presence of an FAD uptake reaction in the model. Finally, the current description of the flavoprotein-related reactions is inconsistent, using sometimes free FAD and sometimes the final electron acceptor. Sometimes both options occur in parallel if the same biochemical reaction occurs twice in the model, due to inconsistencies in metabolite naming. Therefore, the effects of defects associated with FAD biosynthesis or flavoproteins could not, until now, be accounted for by genome-scale constraint-based models. Furthermore FMN is not used as an electron acceptor in the current reconstruction, but only as an intermediate metabolite in the FAD synthesis.

**Fig. 1.**
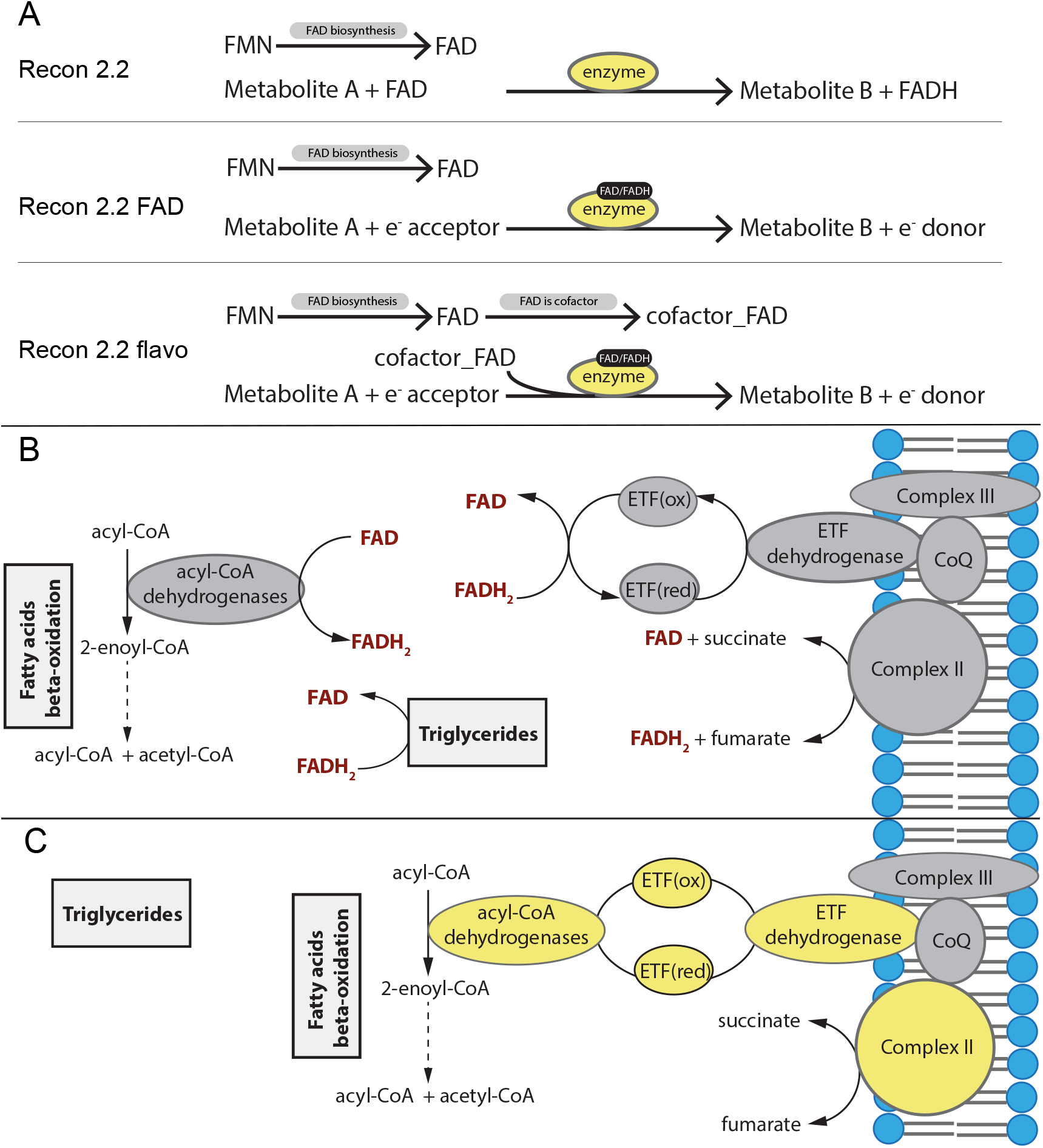
Schematic representation of the difference in handling of the covalently bound cofactors (represented by FAD) between Recon 2.2 and the newly curated models. A. General scheme of the reactions associated with flavoproteins; top: current Recon 2.2; middle: FAD replaced by the final electron acceptor in the reaction equation; bottom: linking the reactions to FAD biosynthesis via the artificial metabolite ‘cofactor_FAD’; B. Scheme of electron transfer from fatty-acid oxidation to the oxidative phosphorylation pathway as in Recon 2.2, C: Scheme of electron transfer from fatty-acid oxidation to the oxidative phosphorylation pathway in our models, Recon 2.2_FAD and Recon 2.2_flavo. Yellow – flavoproteins.

The aim of this study is to explore the physiological effects of flavin-related enzymopathies and to identify candidates for biomarkers. To this end, we modified Recon2.2 to correctly represent flavins as bound cofactors and we introduced a novel simulation approach that enables studies of cofactor scarcity in mammalian models. Moreover, we analysed metabolic disturbances for 38 flavin-related diseases, for which genes were included in the metabolic reconstruction. For 16 of them, which are known to affect the core metabolism, we additionally analysed their ATP production capacity from different carbon sources, showing metabolic blockages in line with the current knowledge about these diseases and their biomarkers.

## 2. Materials and Methods

### 2.1. Genome-scale constraint-based modelling

As described earlier [23] metabolic and transport reactions are summarized in an *m* ×*n* stoichiometric matrix **S**, that contains the stoichiometric coefficient of each of the *m* metabolites in each of the *n* reactions described by the model. Gene–protein-reaction associations use Boolean rules (“and” or “or” relationships) to describe the protein requirement of each reaction.

At steady state, metabolite consumption and production in equilibrium and therefore **b** equals of vector of length *m* with zeros in the following equation:

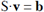

with **v** a vector of length of *n* and vi representing the flux through reaction *i.* Reaction (ir-) reversibility is introduced as constraints limiting minimal and maximal flux values:

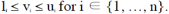

### 2.2. Model curation

We started from Recon 2.2 [3] obtained from the Biomodels database (http://identifiers.org/biomodels.db/MODEL16 03150001). Flavoprotein-related reactions were identified and manually curated based on a review by Lienhart at al. [10] as well as information available in the public databases: KEGG, NCBI Gene and OMIM. Additionally, several rounds of manual curation revealed inconsistencies in directionality of reactions of fatty-acyl CoA ligase, reactions participating in the mitochondrial transport of fatty-acyl carnitines, peroxisomal fatty-acid oxidation and fatty acid synthesis, which were corrected. Missing reactions in the fatty-acid synthesis pathway were added. Subsequently, invalid or duplicated reactions were removed from the model. The curated model was saved as Recon2.2_FAD. For detailed information on all the changes and the underlying documentation, see Supplementary Table 2.

Similarly, we performed a manual curation of FAD-related reactions in Recon 3D model available at https://vmh.uni.lu. We identified all reactions dependent on flavoproteins and manually replaced the FAD and FADH2 with the final electron acceptor (Table S4), creating Recon3D_FAD model.

### 2.3. Model constraints

In simulations of MADD and FAD deficiency, the minimum required flux through the ‘biomass_reaction’ was set to 0.1 mmol•gDW^−1^hr^−1^, to mimic the basic cell maintenance (protein synthesis, DNA and RNA synthesis etc.), unless stated otherwise, as in [25]. Other constraints used only in specific simulations are indicated where applicable. No changes to the model boundaries were made, unless stated otherwise.

### 2.4. MADD simulations

Multiple acyl-dehydrogenase deficiency was simulated as a single gene deletion (ETFDH, HGNC:3483), and the solution space of the models was sampled with the optGpSampler with 10000 points explored and distance of 2000 steps [26]. Additionally, uptake of common carbon_sources (glucose, fructose, C4:0, C6:0, C8:0, C10:0, C12:0 C14:0. C16:0, C18:0, C20:0, C22:0, C24:0, C26:0, alanine, arginine, asparagine, aspartic acid, cysteine, glutamine, glutaric acid, glycine, histidine, isoleucine, lysine, methionine, phenylalanine, proline, serine, threonine, tryptophan, tyrosine, valine, sarcosine) was set to −1 mmol•gDW^−1^hr^−1^

### 2.5 FAD limitation

To study the metabolic response to flavin deficiency we adapted parsimonious FBA [27] to mimic the resource re-allocation among cofactor requiring enzymes upon cofactor scarcity. Furthermore, we linked cofactor requirement and cofactor biogenesis in an approach similar to the method introduced by Beste et al [28]. In short, flavin-related reactions were identified using their GPR annotation (with respect to the Boolean relationship – only reactions for which the flavoprotein is essential were selected), split to two irreversible reactions for each direction and an artificial metabolite ‘cofactor_FAD’ was added to be consumed by each of them with an very low (0.000002) stoichiometric coefficient based on an average human proteome life-time[29] and an average k_cat_ value[30]. Biogenesis of the cofactor was linked to the artificial ‘cofactor_FAD’ through the artificial reaction ‘FADisCofactor’. Adjusting the boundaries of FADisCofactor reaction or reducing the flux of FAD biogenesis pathway allowed us to independently study flavin shortage effects linked to i) FAD biosynthesis impairments, ii) enzymatic mutations hampering cofactor binding or iii) defects in cofactor dissociation and transfer in protein dimers.

Cofactor limitation was obtained by constraining the flux of the FMN adenyltransferase (*FLAT1*) reaction. The solution space for the control model and FAD deficiency model was sampled using optGpSampler with 10000 points explored and distance of 2000 steps [26].

### 2.6 Calculation of maximum ATP yield per carbon source

To calculate the maximum ATP yield per carbon source we adapted the method developed by Swainston et al [3]. All model boundaries for uptake of nutrients were set to 0 except for a set of compounds defined collectively as a minimal medium (Ca^2+^, Cl-, Fe^2+^, Fe^3+^, H+, H_2_O, K+, Na+, NH_4_ SO_4_^2−^, Pi) for which the boundaries of uptake and export fluxes were set to −1000 and 1000 mmol∙gDW^−1^∙h^−1^. Additionally, lower bounds of EX_ribflv(e) were set to −1000 – to simulate riboflavin addition to the minimal medium since the Recon 2.2_flavo model depended on this vitamin to synthesize FAD. For each of the specified carbon sources, the uptake flux was set to −1 mmol∙gDW^−1^∙h^−1^ forcing the model to consume it at a fixed rate. The demand reaction for ATP, ‘DM_atp_c_’ is used to account for cellular maintenance requirements. It was used as an objective function for flux maximization. Under aerobic conditions the possible intake flux of oxygen was set to −1000 mmol∙gDW^−1^∙h^−1^ by changing the lower bound of the corresponding exchange reaction, EX_o2(e).

### 2.7 Analysis of biomarkers for inborn errors of metabolism

Inborn errors of metabolism and known genes mutated in these IEMs were retrieved from the OMIM database (34, https://www.omim.org). GPR associations in the model were used to identify affected reactions and they were subsequently removed from the model for the simulation of the corresponding disease. Maximum ATP yield was calculated for a set of carbon sources linked to known biomarkers, as described above. Changes in the maximum ATP yields between the diseased model and its healthy counterpart were compared to reported biomarkers [32].

Secondly, a method described originally by Shlomi et al. [24] and modified by Sahoo et al. [33] was used to screen for potential biomarkers. In short, the min and max values of each exchange reaction were calculated in disease or healthy models. In the disease models a gene knock-out was simulated by simultaneously blocking all reactions associated with the tested gene. In the healthy models all reactions associated with the tested gene were simultaneously activated. Min-max intervals were then compared to test if the flux range would yield a higher or lower exchange reaction flux, and as the result, would lead to increased or decreased concentrations of a studied metabolite in the bodily fluids (blood/plasma, urine).

A list of tested single enzyme deficiencies together with their respective objective functions and biomarkers has been provided (See Table S3 [Known biomarkers & diseases]).

### 2.8 Software

Model curation and all simulations were carried out with MatLab R2015a (MathWorks Inc., Natick, MA) using the Gurobi5.6 (Gurobi Optimization Inc., Houston TX) linear programming solver and the COBRA toolbox [34]. For FVA analysis GLPK 4.63 (GNU Linear Programming Kit, https://www.gnu.org/software/glpk/) solver was used. Data analysis was performed using MatLab R2015a. All supplementary files, models, and scripts can be found at https://github.com/WegrzynAB/Papers.

## 3. Results

### 3.2 A new representation of flavoprotein biochemistry

We started by manually curating the existing Recon 2.2 model to improve the simulation of flavin dependent reactions. We assembled a list of 111 flavoproteins (Table S1 [Flavoproteins]). The genes encoding 62 of these proteins were already present in Recon 2.2 and they were associated with 378 reactions. First, the gene-protein-reaction (GPR) associations in the selected reactions were examined. Wrongly annotated genes were corrected. In the mitochondrial and peroxisomal beta-oxidation pathways, many GPR associations were extended to account for complete oxidation of fatty acids. Secondly, the directionalities of fatty-acyl CoA ligases, reactions participating in the mitochondrial transport of fatty-acid-carnitines, and fatty acid synthesis reactions were checked, and inconsistencies were corrected. Lastly, missing reactions in unsaturated fatty acid synthesis and peroxisomal beta-oxidation were added. Altogether 548 reactions were corrected, 31 were added, and 49 reactions were deleted from the model (Table S2). In the original Recon 2.2 model FAD was represented as a free metabolite similar to NADH (Fig 1A, top). This approach artificially generated a pool of free FAD, while physiologically FAD is a part of an active holoenzyme rather than a free metabolite. Thus, systemic effects of flavoprotein deficiencies could not be properly accounted for. We have changed the model to more precisely describe the cofactor dependency of these reactions. To this end, FAD has first been removed as a free metabolite and replaced by the final electron acceptor in each reaction, such as ubiquinone-10 or ETF (Fig. 1A, middle). This approach allows a more accurate representation of the electron channelling in the pathway, which is clearly visible in the scheme describing how fatty-acid oxidation links to oxidative phosphorylation (Fig 1C). At this stage, 157 reactions were curated and 33 reactions were deleted. This curated model version was called Recon 2.2_FAD.

Removing FAD and FMN as free metabolites from all redox reactions, however, made these reactions completely independent from flavin biosynthesis. To preserve a link between riboflavin dependency and flavoprotein-related reactions, an artificial metabolite called ‘cofactor_FAD’ was generated from the newly synthesized FAD (Recon2.2_flavo model). This cofactor was added as an additional substrate to all flavoprotein-dependent reactions, to be consumed with a low stoichiometric coefficient (Fig. 1A, bottom). The latter was estimated based on available data on protein half-lives [29] and catalytic turnover rates (kcat) [30] in humans. Thus, it can be seen as a flavin-maintenance requirement. In the resulting model version all flavoprotein-dependent reactions are strictly dependent on the presence of riboflavin and flavin biosynthesis, yet without introducing the artificial degrees of freedom of the Recon 2.2 model. We have additionally tested if the chosen stoichiometric coefficient did not overly constrain the model. To this end we tested the sensitivity of the flavoprotein-related reaction flux through the model to changes in the cofactor stoichiometric coefficient. Only a minor effect on the flux was observed (Fig. S4B). In total 420 reactions were linked to the FAD biosynthesis by cofactor consumption.

### 3.3 Metabolic role of the flavoproteome

We mapped the known human flavoproteins (Table S1) to the original and the updated versions of Recon 2.2. For each flavoprotein we identified all the associated metabolic reaction(s). Also reactions for which a non-flavoprotein isoenzyme was present, were included in the mapping. We found that the majority (270 out of 381 for Recon 2.2 and 263 out of 401 for Recon 2.2_FAD/flavo) of the reactions associated with flavoproteins were localized in peroxisomes and mitochondria (Fig. S1A). Out of the 65 flavoprotein genes included in the Recon 2.2 38 are linked to known human diseases (Table S1 [Diseases]). Moreover, our curation added seven additional disease-linked flavoproteins genes (*NOS1, MTRR, NQO2, L2HGDH, IYD1*, *D2HGDH* and *FOXRED1*) to the Recon 2.2 gene set (Fig. 2A). In total 73% of the disease-linked flavoproteins were mapped onto Recon 2.2_FAD/flavo, while 69% of them were mapped on the Recon 2.2 model. Among the disease-linked flavoproteins that were not included in our models are those of non-metabolic function: a chaperone (*ALR*), a pro-apoptotic factor (*AIFM1*), a histone demethylase (*KDM1A*), and a proton channel (*NOX1*).

**Fig. 2.**
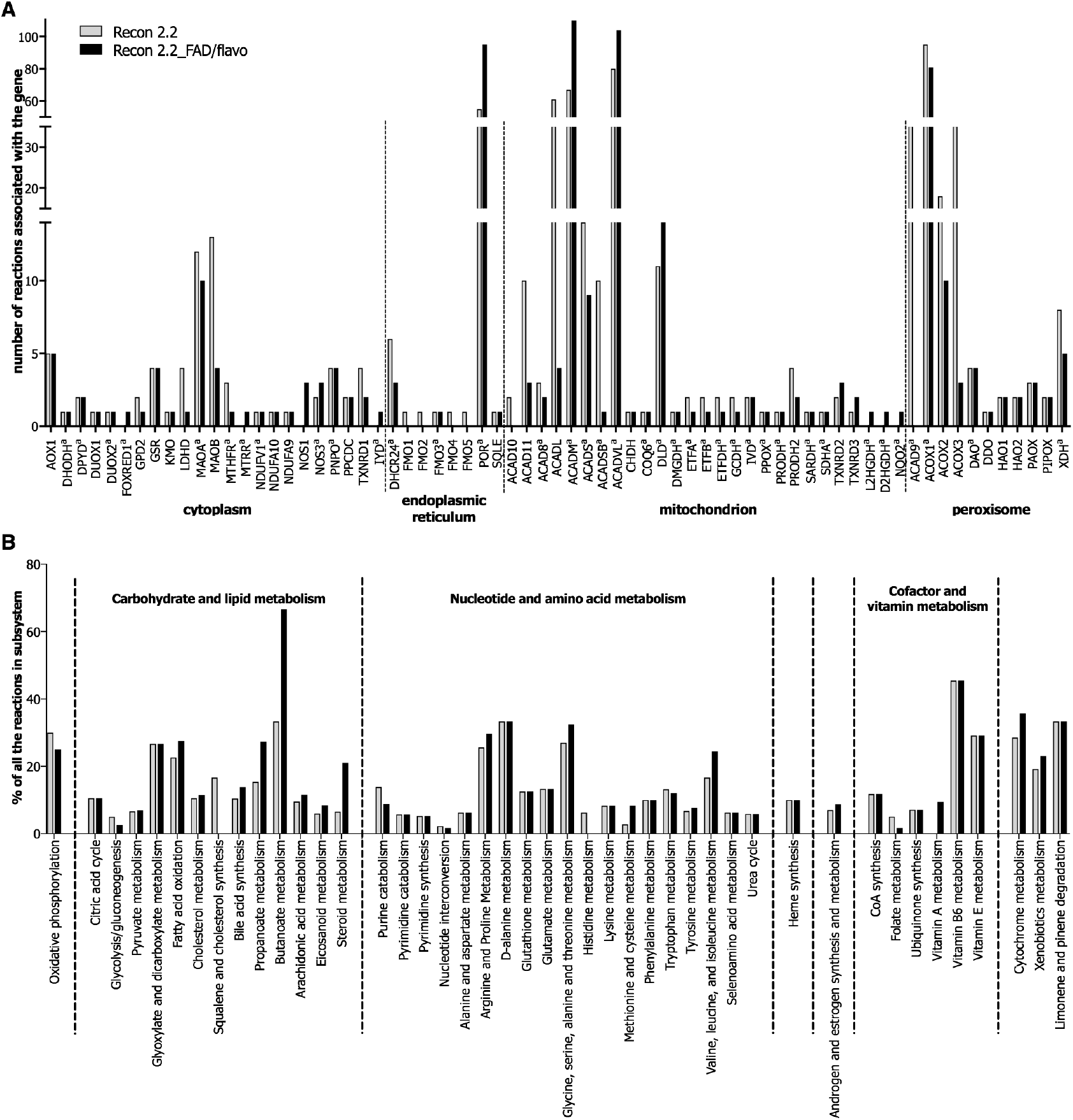
Mapping of the flavoproteome on the models before (grey, Recon 2.2) and after curation (black, Recon 2.2_FAD or Recon 2.2_flavo). A. Number of reactions associated with each flavoprotein-encoding gene. amarks a gene associated with an IEM; B. Percentage of reactions associated with flavoproteins per subsystem (subsystems as defined in Recon 2.2).

We then investigated the reliance of metabolic subsystems, as defined in the original Recon 2.2 model, on flavoproteins. Vitamin B6 metabolism, cytochrome metabolism, butanoate metabolism, D-alanine metabolism, and limonene and pinene detoxyfication were most heavily affected with more than 30% of their reactions depending on flavoproteins (Fig. 2B). Additionally, oxidative phosphorylation was also affected with 2 out of 8 of its reactions linked to the flavoproteins (Fig. S1B).

### 3.4 The improved model correctly simulates MADD

To simulate MADD, we deleted the *ETFDH* gene (HGNC:3483; Fig 1B and C) from all model versions and calculated the steady-state solution space. Indeed, the flux through the ETF dehydrogenase reaction was completely blocked in all three models, confirming that the deletion was effective (Fig. S2A). The biomass production flux remained largely unchanged in all the models (Fig. S2B). This is not surprising, since the simulations were performed with carbon and energy sources other than fatty acids available, such as sugars and amino acids. As we had suspected, deletion of *ETFDH* did not dramatically reduce the fatty-acid oxidation flux in the original Recon2.2 model (Fig. 3).

**Fig. 3.**
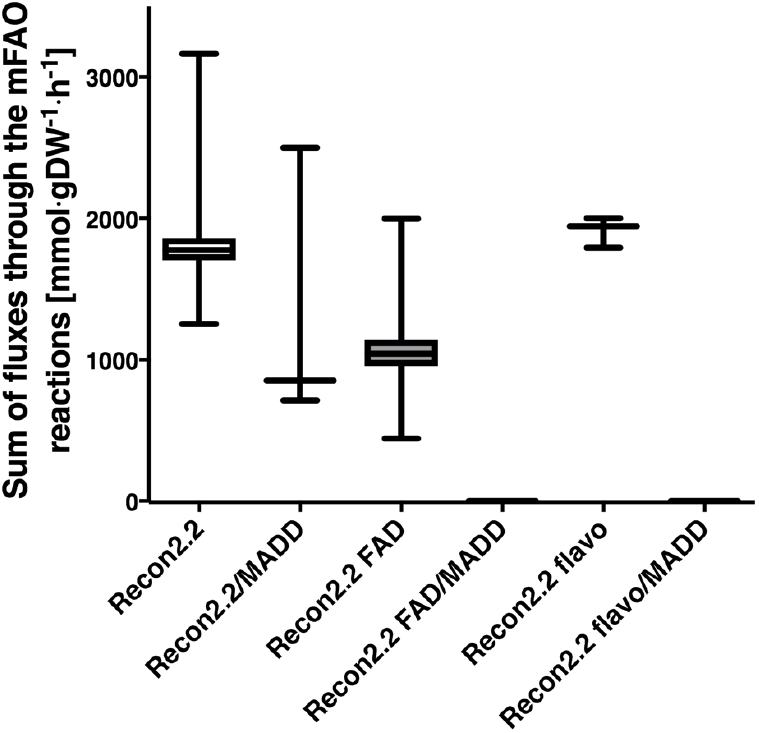
New models can correctly simulate the physiology of MADD. Overall flux through the mFAO (sum of fluxes through the mFAO reactions). Only in the new models Recon2.2_FAD and Recon2.2_flavo full block of the mitochondrial fatty-acid oxidation flux was observed upon deletion of *ETFDH*. Flux values were obtained by sampling the solution space using the optGpSampler (10000 points explored, 2000 steps). Boxes span 25^th^ to 75^th^ percentiles, lines show medians, whiskers indicate minimum and maximum values.

This can be explained from the fact that in Recon 2.2 the FADH2 produced by the acyl-CoA dehydrogenases in the fatty-acid beta-oxidation is not strictly coupled to ETF dehydrogenase, but can be reoxidized by other reactions utilising FADH2 (Fig. 1B). This prediction is clearly incorrect, since mitochondrial fatty-acid oxidation is severely impaired in MADD patients [13]. In contrast, deletion of *ETFDH* caused a total block of the mitochondrial fatty-acid oxidation in Recon2.2_FAD and Recon2.2_flavo (Fig. 3), in agreement with the disease phenotype. Since we had, as part of the curation, removed the incorrect FAD reaction from the model, we checked that this could not explain our results. Indeed, the results were not altered by addition of free FAD uptake reaction back to the Recon2.2_FAD model, neither by only deleting free FAD uptake reaction from Recon 2.2 model (Fig. S2C).

Subsequently we compared the maximum stoichiometric ATP yield per unit of consumed carbon source in aerobic conditions between the models. If a substrate was incompletely metabolized, this caused a decrease of the maximum ATP yield. If the substrate could not be metabolised to produce ATP, then the obtained yield was zero. In contrast to the original model, Recon2.2_FAD and Recon2.2_flavo predicted that *ETFDH* deletion eliminated any ATP yield from fatty-acids with chain lengths C*4* through C*14*, which are primarily substrates of mFAO in the models. Longer fatty acids (C*16* to C*26*) can be partially oxidized by peroxisomal beta-oxidation and therefore still yield some ATP if *ETFDH* is deficient, also in Recon2.2_FAD and Recon2.2_flavo (Table 1). Instead of the mitochondrial acyl-CoA dehydrogenase, the peroxisomal pathway contains a hydrogen peroxide producing acyl-CoA oxidase, which is not dependent on the ETF system as electron acceptor. Glycolysis and the TCA cycle remained unchanged, since they do not depend on *ETFDH* either. Accordingly, aerobic sugar metabolism was not affected by *ETFDH* deficiency in any of the models (Table 1).

**Table 1.** Comparison of maximum ATP yields in a medium without riboflavin (B2 -), with riboflavin (B2 +), and in a model with *ETFDH* knocked out in the presence of B2 (MADD). Minimal medium: Ca^2+^, Cl^−^, Fe^2+^, Fe^3+^, H^+^, H_2_O, K^+^, Na^+^, NH_4_ SO_4_^2−^, Pi, plus the indicated carbon and energy source. Note that this medium is insufficient for cell growth. ATP yield was expressed stoichiometrically as the net rate of ATP production (in steady-state equal to the rate of the ATP demand reaction) divided by the rate of carbon-source consumption in mmol ∙ g DW^−1^ ∙ h^−1^. If no ATP was produced at all, the yield was set to zero, irrespective of whether the carbon source was consumed, to avoid division by zero. We verified that the precise value of the stoichiometric coefficient for FAD consumption in Recon2.2_flavo did not affect the maximum ATP yield from single carbon sources at the accuracy presented here.

| Carbon source | ATP yield |  |  |  |  |  |  |  |  |  |
| --- | --- | --- | --- | --- | --- | --- | --- | --- | --- | --- |
|  | Theoretical | Recon2.2 |  |  | Recon2.2_FAD |  |  | Recon2.2_flavo |  |  |
|  |  | B2 - | B2 + | MADD | B2 - | B2 + | MADD | B2 - | B2 + | MADD |
| Glucose | 31 | 32 | 32 | 32 | 32 | 32 | 32 | 2 | 32 | 32 |
| Fructose | 31 | 32 | 32 | 32 | 32 | 32 | 32 | 2 | 32 | 32 |
| C4:0 | 21.5 | 22 | 22 | 22 | 22 | 22 | 0 | 0 | 22 | 0 |
| C6:0 | 35.25 | 36 | 36 | 36 | 36 | 36 | 0 | 0 | 36 | 0 |
| C8:0 | 49 | 50 | 50 | 50 | 50 | 50 | 0 | 0 | 50 | 0 |
| C10:0 | 62.75 | 64 | 64 | 64 | 64 | 64 | 0 | 0 | 63.99 | 0 |
| C12:0 | 76.5 | 82.5 | 82.5 | 82.5 | 78 | 78 | 0 | 0 | 77.99 | 0 |
| C14:0 | 90.25 | 92 | 92 | 92 | 92.5 | 92.5 | 0 | 0 | 92.49 | 0 |
| C16:0 | 104 | 106.75 | 106.75 | 106.75 | 108.25 | 108.25 | 2.25 | 0 | 108.24 | 0 |
| C18:0 | 117.75 | 120 | 120 | 120 | 121.5 | 121.5 | 12.75 | 0 | 121.49 | 12.75 |
| C20:0 | 131.5 | 134 | 134 | 134 | 135.5 | 135.5 | 25 | 0 | 135.49 | 25 |
| C22:0 | 145.25 | 147.25 | 147.25 | 147.25 | 147 | 147 | 38 | 0 | 146.99 | 38 |
| C24:0 | 159 | 160.5 | 160.5 | 160.5 | 159.25 | 159.25 | 50.25 | 0 | 159.24 | 50.25 |
| C26:0 | 172.75 | 170.75 | 170.75 | 170.75 | 169.5 | 169.5 | 61.75 | 0 | 169.49 | 61.75 |

Cn:0, a saturated fatty acid of length n
<sup>a</sup> Theoretical yields as calculated by Swainston et al. after Salway[50]

### 3.5 FAD deficiency

Subsequently, we studied the systemic effects of riboflavin deficiency caused by reduced FMN and FAD synthesis from their precursor riboflavin (vitamin B2). Mutations of FAD synthase (encoded by the *FLAD1* gene) are known to exist. Some mutants maintain residual catalytic activity and are therefore responsive to dietary riboflavin, while others are completely non-responsive to riboflavin supplementation [35]. *FLAD1* mutations cause MADD-like symptoms in patients, with elevated levels of multiple acylcarnitines and organic acids in blood and/or urine [35].

We repeated the calculation of ATP yields per unit of carbon source with and without riboflavin. In Recon2.2_FAD model, FAD had been removed as an electron acceptor from all flavoprotein-dependent reactions, thus completely uncoupling these reactions from FAD biosynthesis (Fig. 1A). Accordingly, the ATP yield of none of the substrates was affected by the absence of riboflavin in this model (Table 1). In contrast, in Recon 2.2_flavo all flavoprotein-dependent reactions are strictly dependent on the presence of flavins (Fig. 1A). Therefore, if FAD and FMN are depleted due to riboflavin deficiency, none of the flavoprotein-dependent reactions can run anymore. Consistently, we found that in the Recon 2.2_flavo model the mitochondrial beta-oxidation and the TCA cycle were fully blocked in the absence of riboflavin. Therefore, ATP could only be generated in glycolysis, leading to only 2 ATP per sugar molecule and no ATP was generated from any of the fatty-acid substrates in the absence of riboflavin. We noted that the net ATP yields of all substrates in the presence of riboflavin were lower in Recon 2.2_flavo than in Recon2.2_FAD (Table 2). The reason is that the active FAD biosynthesis in Recon2.2_flavo requires ATP.

**Table 2.** Comparison of the prediction accuracy of Recon 2.04, Recon 2.2 and Recon 2.2_flavo against the gold standard (GS) [17] or against our compendium of flavoprotein-related diseases (FD).

|  | Recon2 | Recon2.2 | Recon2.2_flavo | Recon2.2 | Recon2.2_flavo |
| --- | --- | --- | --- | --- | --- |
|  | GS | GS | GS | FD<sup>a</sup> | FD |
| Correct prediction | 76 | 88 | 94 | 11 | 16 |
| Wrong prediction - wrong direction<sup>a</sup> | 23 | 29 | 30 | 3 | 11 |
| Accuracy<sup>a</sup> | 77% | 75% | 76% | 78% | 59% |
| P value (Fischer test) | $7.93 \times 10^{-4}$ | $1.2 \times 10^{-2}$ | $2.6 \times 10^{-3}$ | 0.16 | 0.12 |
| Wrong prediction – no change<sup>a</sup> | 187 | 169 | 162 | 53 | 39 |
| True positive rate<sup>a</sup> | 26% | 31% | 33% | 17% | 24% |

<sup>a</sup> wrong direction – where the model predicted the metabolite to change in the opposite direction from the experimental values
- <sup>b</sup> Prediction accuracy measured as a ratio of correct positive predictions to sum of all predicted biomarkers (correct and wrong direction) as reported in Thiele et al.[23] <sup>c</sup> no change - where model predicted no change in the metabolite, despite it being known as a biomarker <sup>d</sup> True Positive Rate measured as a ratio of correct predictions to the sum of all biomarkers in the database.

Finally, we analysed the response of the models to a gradual limitation of the rate of the FAD synthase reaction (Fig. 4A). First, we computed how the maximum flux capacity of the summed flavoprotein-catalysed reactions responded to a limitation of FAD biosynthesis. The Recon 2.2_FAD model remained unresponsive to the rate of FAD biosynthesis (Fig. 4B). This is not surprising, since in Recon 2.2_FAD, the final electron acceptor replaced FAD in each reaction; hence, all flavoproteins were artificially independent of the FAD. Recon 2.2_flavo instead made the flavoprotein-catalysed reactions dependent on a low rate of biosynthesis (Fig. 1A). This resulted in a linear decrease of the flavoproteome-dependent flux capacity in Recon2.2_flavo, when the FAD synthesis flux was below 0.05 mmol×gDW^−1^×h^−1^. Subsequently, we calculated how the shape of the actual steady-state solution space for the FAD-related reactions changed due to the cofactor_FAD limitation. The overall average steady-state flux (Fig. 4C) was lower than its maximum capacity (Fig. 4B). The responsiveness of the Recon 2.2_flavo model was preserved but since the pathways worked below their maximum capacity, the cofactor_FAD limitation started to play a role only when the cofactor availability was close to zero (Fig. 4C).

**Fig. 4.**
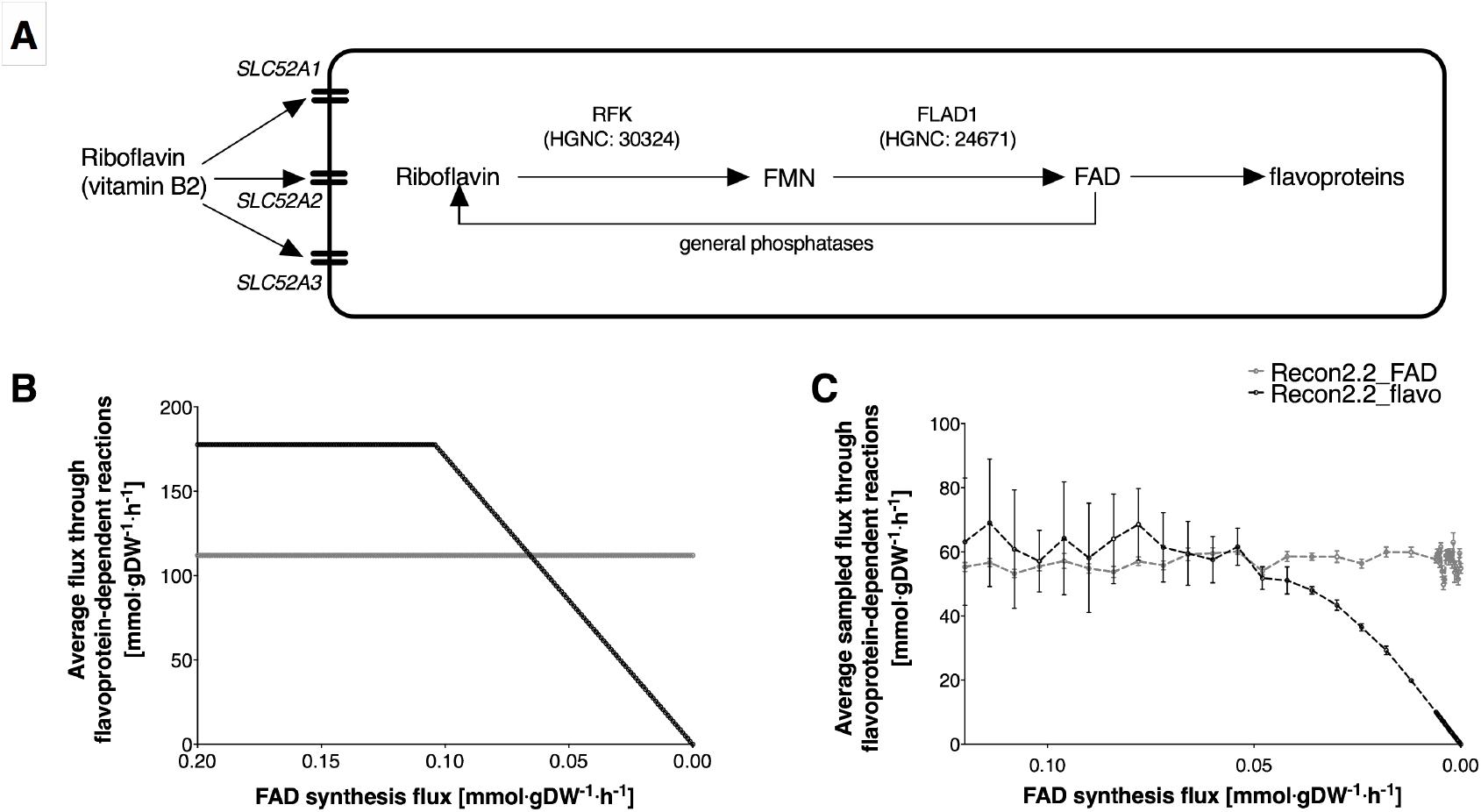
Coupling of FAD-related reactions to FAD-biosynthesis enabled the new model to respond to low cofactor availability. A. Schematic representation of FAD biosynthesis, *RFK* – riboflavin kinase, *FLAD1* flavin adenine dinucleotide synthetase 1. B. Average flux through the FAD-related reactions as a function of FAD biosynthesis flux. Maximum capacity was calculated by maximizing the objective function – a sum of fluxes through all the FAD-related reactions. C. Average and SD of the flux through the FAD-dependent reactions from the sampled solution space (using the optGpSampler with 10000 points explored and 2000 steps distance) in response to declining cofactor biosynthesis flux.

### 3.6 Prediction of biomarkers

Genome-scale metabolic-models have been shown to assist in biomarker prediction. The authors of the original Recon 2.04 model [23], used the method by Sahoo et al [33] to predict biomarkers based on altered maximum uptake and secretion rates of metabolites. We tested the same method for biomarker prediction in Recon 2.04, Recon2.2 and Recon2.2_flavo models against the golden standard set of known biomarkers [17]. Prediction accuracies (correctly predicted biomarkers compared to total predicted biomarkers), and their p-values, calculated as reported in Thiele et al. [23] were similar in all the models (Table 2, GS) and in agreement with the previously published 77% accuracy in Recon 2 [23]. With respect to True Positive Rate (the percentage of known biomarkers that was correctly predicted) our Recon 2.2_flavo model scored highest with 33%, while Recon 2.2 and Recon 2.04 had 31% and 26% TPR respectively (Table 2). This shows that we could recover more biomarkers without a loss in accuracy. Furthermore, we used the same method to study flavoproteome-related diseases and their associated biomarkers (Table S3 [Known biomarkers&diseases]) in our Recon 2.2_flavo model. For this subset of diseases TPR was only 24%, coupled with lower accuracy of 59%. The p-value of the prediction accuracy was higher for this subset of diseases (Table 2, FD). The high p-value can be linked to the relatively small number of diseases in our FD set and therefore a higher chance of obtaining the same values distribution by chance. The low TPR values for all tested models reflect the FVA-based method’s inability to predict many known biomarkers for important flavoproteome-related diseases, such as those in fatty-acid metabolism and oxidative phosphorylation (Table S3 [FVA]). For instance, typical biomarkers of fatty-acid oxidation defects, acyl carnitines were not altered in disease simulations.

To circumvent this caveat of the classical method, we decided to study the maximum ATP yields for a selected range of carbon sources that are also biomarkers in human diseases, modifying the method of Swainston *et al.*[3]. This method allows to study the capacity to metabolise a carbon source under defined model boundaries and compare the outcome from the disease model and control model. We selected 16 diseases which are known to affect core metabolism and for which we expected changes in the net ATP production from certain carbon sources. In total for 11 out of 16 diseases studied, the known biomarkers, marked in red boxes in Fig. 5, could be linked to the metabolic changes seen in the Recon 2.2_flavo model. For *FOXRED1*, *PRODH* and *ETFDH* deficiencies the Recon 2.2 model did not predict any metabolic changes while our flavoproteome-curated models showed reduced ATP yield from carbon sources known as biomarkers for these diseases. Moreover, our method revealed disrupted amino acid metabolism in *ACADSB, DLD, GCDH*, *IVD*, and *ACAD8* deficiencies, which may point to new possible biomarker profiles. In all MADD variants (*ETFDH*, *ETFA* or *ETFB* deficiency) a full block of mitochondrial FAO capacity was seen only in our improved models. Long-chain fatty acids (above C20:0) were still partially metabolised as peroxisomal FAO remained functional. In contrast, and as expected, when simulating deficiency of the peroxisomal enzyme *ACOX1*, only the peroxisomal FAO was impaired.

**Fig. 5.**
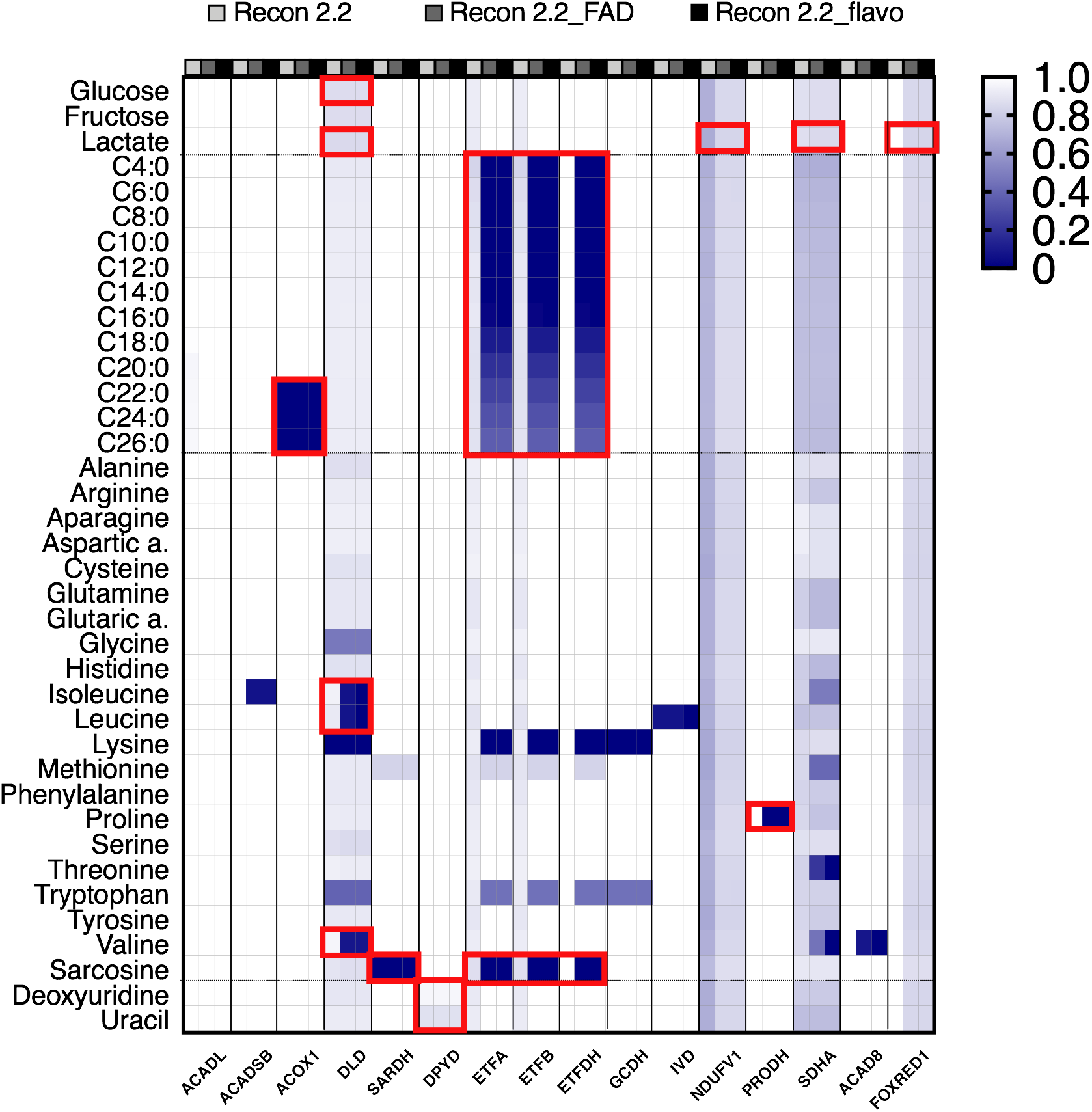
Changes in the ATP yield from different carbon sources in flavoprotein-related diseases predict metabolic adaptations in energy metabolism. For each disease, predictions from all three models are shown, first (light-grey): Recon 2.2, second (grey): Recon 2.2_FAD, third (black): Recon 2.2_flavo. Ratios between ATP yields in disease model vs. healthy model are shown as a gradient (white – no change (1), dark blue – reaction is blocked in disease model (0)). Red frames - metabolites known to be affected, elevated in blood and/or plasma, in patients. See Supplementary Table 3 [Known biomarkers&diseases] for details about the IEM’s and the reactions used as biomarkers. *ACADL* – Acyl-CoA dehydrogenase very long-chain deficiency (MIM:201475); *ACADSB* - Short/branched-chain acyl-CoA dehydrogenase deficiency (MIM:610006); *ACOX1* – Adrenoleukodystrophy (MIM:264470); *DLD* – Dihydrolipoamide dehydrogenase deficiency (MIM:246900); *SARDH*– sarcosinemia (OMIM entry MIM:268900); *DPYD* - Dihydropyrimidine dehydrogenase deficiency (MIM:274270); *ETFA* - Glutaric aciduria IIA (MIM:231680); *ETFB* - Glutaric aciduria IIB (MIM:231680); *ETFDH* - Glutaric aciduria IIC(MIM:231680); *GCDH*– glutaryl-CoA dehydrogenase deficiency (MIM:231670); *IVD*– Isovaleric academia (MIM:243500); *NDUFV1* - Leigh syndrome (MIM:256000); *PRODH* - Hyperprolinemia 1 (MIM:239500); *SDHA* - Mitochondrial complex II deficiency (MIM:252011); *ACAD8* - Isobutyryl-CoA dehydrogenase deficiency (MIM:611283); *FOXRED1*– mitochondrial complex I deficiency (MIM: 252010);

For many diseases, metabolic biomarkers are not known, or the known biomarkers fail to differentiate between patients with different severity of the disease. Our models predicted metabolic changes at the level of maximum ATP yield per carbon source in all tested flavin-related diseases. Clear disease-specific patterns are seen, which allow to distinguish between different diseases. Most of the changes were identified in the amino acids metabolism. Those compounds are routinely used in the newborn screening test; however, they have not yet been related to these diseases.

### 3.7 Compatibility with Recon 3D

During the writing of this manuscript Recon 3D came out [4], another major update of the human metabolic reconstruction that had been developed in parallel to Recon 2.2 [3]. The number of reactions was increased by 74% to 13,543, and the number of metabolites by 56% to 4,140 in Recon 3D compared to Recon 2.2. Furthermore, the representation of enzymes involved in oxidative phosphorylation system has been corrected according to the fix made in Recon 2.2 [3]. However, the change of the gene mapping from GeneID to HGNC standard proposed in Recon 2.2 was not applied in Recon 3D. We tested the applicability of our curation to Recon 3D, in order to ensure that it can be readily used with different model versions. Similar to previous model versions, Recon 3D treats FAD as a free cofactor. Our method of replacing FAD with the final electron acceptor was easily applied to Recon 3D (Table S4). The impact of FAD cofactor limitation was similar as in the Recon2.2-derived models (Fig. S4A). Additionally, we tested how Recon 3D responded to simulation of MADD. As seen before, there was a significant remaining total flux through the mFAO reactions in the Recon 3D model with *ETFDH* deficiency, while in the flavoproteome-curated models the mFAO flux was correctly blocked by MADD (Fig.S5A). We verified that the deletion was effective (Fig. S5B), and that the biomass production flux remained largely unchanged in all the models (Fig. S5C).

Since Recon 3D has a much larger metabolic coverage than its predecessors, we tested its capability of detecting biomarkers. However, the model showed no substantial change in the accuracy (74%) of biomarker predictions, while a significant decrease in the TPR value (10%) was observed (Table S5).

## 4. Discussion

We present an updated version of Recon 2.2 that was curated and extended to correctly represent the flavoprotein-catalysed reactions. Furthermore, we introduced a new method to study the role of enzyme-bound cofactors, such as FAD. Curating the representation of FAD in Recon 2.2 allowed to correctly simulate aberrant metabolic behaviour upon single enzyme deficiencies. Since the predecessor of Recon 2, the metabolic reconstruction Recon 1 [36], was published, many groups have extended and improved model versions. They used it as a basis for tissue specific models [5,23,37–40], studied the effects of diet [38], and predicted biomarkers for enzymopathies [17,23,24,38]. Work by Smallbone [41] and Swainston et al. [3] focused on a full mass and charge balance and on simulations of energy metabolism. However, despite their crucial role in metabolism, none of the curative efforts in human reconstructions, including the most recent Recon 3D [4], focused on cofactors, not even organic cofactors that are (in part) synthesized in the cell. Metabolism related to other apoenzymes requiring other bound cofactors for their activity (metals, iron-sulfur clusters, or heme) would potentially profit from the same solution to further enhance biomarker research in genome-scale models.

Flavoprotein-linked diseases can lead to very strong metabolic responses in patients, such as episodes of severe metabolic derangement, hypoglycaemia, metabolic acidosis, sarcosinemia and cardiovascular failure in MADD patients. Acylcarnitines, as well as sarcosine are known to be changed in the plasma and urine of MADD patients [12–14]. The original Recon 2.2 model could not predict any of the known biomarkers and no systemic effects of MADD were seen in the simulations, because the model incorrectly comprised alternative routes to reoxidize FADH2. In our new model, in which the electrons are transferred to the final electron acceptor of each flavoprotein-catalysed reaction rather than to a soluble FAD pool, and in which flavoprotein-dependent reactions are dependent on flavin synthesis, both systemic effects and metabolic changes linked to biomarkers were predicted correctly. This is seen clearly by a full block of mFAO capacity while peroxisomal FAO remained functional in MADD. In contrast, when simulating deficiency of the peroxisomal enzyme ACOX1, only the peroxisomal FAO was impaired, leading to reduced metabolism of long-chain fatty acid substrates (Fig. 5). This extension is relevant for a correct description of mitochondrial fatty-acid oxidation defects, which can be partly rescued by peroxisomes [42].

In total, we tested metabolic changes for 45 diseases, out of which 31 are associated with biochemical biomarkers. A caveat of the existing methods for biomarker predictions is that they only include the metabolites that are known as biomarkers. The models, however predict many more metabolites with altered production or consumption rates. These are potential novel biomarkers. Since they have most often not been explored experimentally, however, we do not know if the predictions are correct. If these would be tested, we would get a more complete insight into the accuracy of our predictions. Therefore, we propose usage of true positive rates for more correct description of model performance. Using Recon 2, Recon 2.2, and Recon 2.2_flavo, we predicted biomarkers for diseases included in the compendium of inborn errors of metabolism published by Sahoo et al. [17] with True Positive Rates of 26%, 31% and 33% respectively, while accuracies, as calculated in Thiele et al. [23], remained similar to previously published 77% (77%, 75% and 76% respectively). A lower (17% and 24%) TPR was reported with Recon2.2 and Recon2.2_flavo respectively for biomarkers of the flavoprotein-related diseases subset, with accuracies of 78% and 59% respectively. However more detailed studies of metabolism, using ATP production yield estimation performed for 16 flavoprotein-related diseases linked to the core metabolism, showed promising results for both our models. This method allowed us to test if alternative metabolic pathways exists that allow ATP production from the single carbon sources in various IEMs. The metabolic changes identified with this method were in line with clinical data, including impaired FAO and sarcosine degradation in all MADD cases, no proline degradation in *PRODH* deficiency and blocked very-long chain FAO in *ACOX1* deficiency [43]. Interestingly, ATP-generating breakdown of amino acids has been predicted to be affected in several diseases analysed. Valine breakdown has been predicted by our model to be significantly impaired in isobutyryl-CoA dehydrogenase deficiency (*ACAD8)* which is in line with the literature knowledge about this disease [44]. Furthermore, our models predict a decreased ATP yield from breakdown of several amino acids, and a general impairment of energy metabolism in *SDHA* deficiency. This extremely rare disease is indeed known to affect energy metabolism. However, due to its low prevalence, no specific biomarkers are known [45]. Our models predict valine, leucine, threonine, and methionine degradation pathways to be most severely affected in this disease. FAD-containing enzymes are crucial in both fatty acid oxidation and amino acid metabolism as was highlighted in flavoproteome mapping (Fig. 2B and S1B). Consistently, their impact on biomarkers became more pronounced after our curation (Fig. 5). For all 16 tested flavoprotein– related diseases we predicted new metabolic changes that may lead to new biomarker patterns. Our data suggests that these diseases might have multiple identifiable biomarkers. Using a multimarker approach or specific biomarkers ratios, which is common in cardiovascular risk assessment [46], instead of only single compounds, we could better differentiate between different diseases and potentially also between patients with different severity of the defects which has been proven recently for Zellweger syndrome patients differentiation [19]. The latter was not pursued here, since we only studied complete enzyme deficiencies.

Limitations in accuracy and True Positive Rates of biomarker predictions with existing methods and models, may have different causes. Cellular lipid profiles are very complex [19], and currently incompletely represented in the human genome-scale models. Since many flavoprotein-related diseases affect lipid metabolism it is likely that a better representation of the lipid metabolism will improve the predictions. Additionally, our new method of cofactor implementation could be extended to account for all different cofactors required in human metabolism. Future improvement of our method may involve a differentiation in stoichiometric coefficients of flavin usage per enzyme depending on the specific protein half-life. This would allow the incorporation of differences in efficiency of flavin utilization for various flavin-dependent enzymes, thereby increasing the accuracy of metabolic predictions. In addition, one should remain critical on our assumption that all FAD or FMN is tightly bound as a prosthetic group. While some flavoproteins have FAD covalently bound, most have a non-covalent, yet tight-binding FAD or FMN. Some flavoproteins, however, may have a relatively low FAD binding affinity. This holds for instance for bacterial two-component monooxygenases in which reduced FAD must be translocated from one protein domain to another [47]. Low FAD affinity of cancer-associated variants of NAD(P)H quinone oxidoreductase 1 leads to low protein stability [48]. We are not aware of low-affinity flavoproteins that depend on free diffusion of reduced FAD.

We noted that the currently most extensive reconstruction of human metabolism, Recon 3D, showed a significant decline in the number of correctly predicted biomarkers compared to its predecessors (Table S5). By extending the coverage of the metabolic network, alternative pathways have been created. One may hypothesise that their physiological relevance is smaller in reality than in the model, e.g. due to kinetics, spatial separation, or thermodynamics. This limitation can possibly be overcome by using tissue-specific models with an appropriate set of boundaries for the exchange reactions, as has been proposed recently by Thiele et al. [49]. Finally, it is quite likely that some biomarkers will only be predicted correctly when kinetic and thermodynamic constraints are included.

## Acknowledgements

We would like to thank prof. D.J. Reijngoud for valuable discussions, particularly regarding the interpretation of biomarker predictions.

This work was supported by the Marie Curie Initial Training Networks (ITN) action PerFuMe [project number 316723] and the University Medical Center Groningen. BMB was further supported by a CSBR grant from the Netherland Organization for Scientific Research (NWO) supporting the Systems Biology Centre for Energy Metabolism and Ageing [853.00.110].

## Supporting information

**Fig S1.**
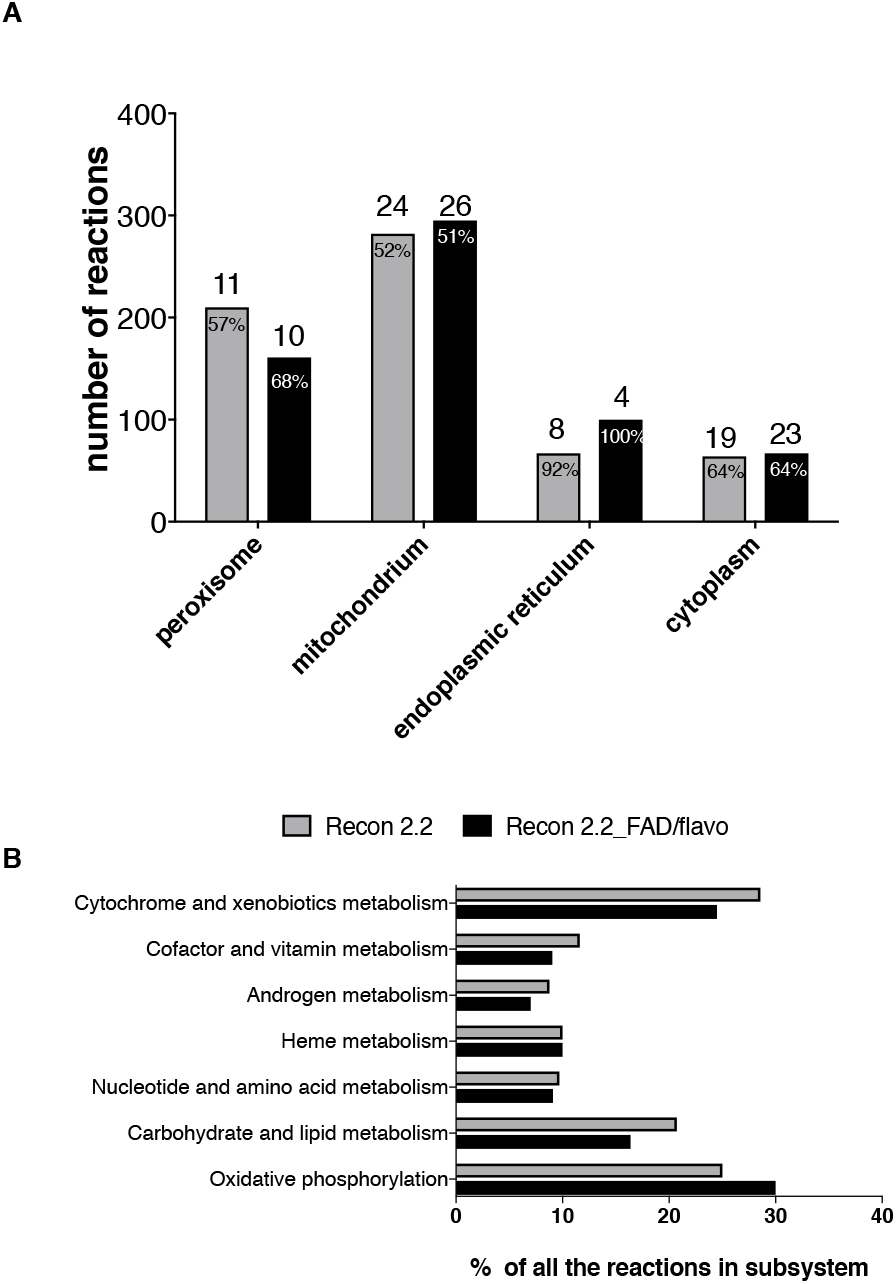
Mapping of the flavoproteome on the models before (grey, Recon 2.2) and after curation (black, Recon 2.2_FAD/flavo). A. Number of flavoprotein-related reactions in each compartment. Numbers above the bars show the number of flavoproteins in the compartment. Values in the bars show the percentage of all reactions in the subsystem that are affected by flavoproteins. B. Average percentage of the reactions associated with flavoproteome in the subsystem grouped by higher level metabolic functions (as defined in Recon 2.2).

**Fig S2.**
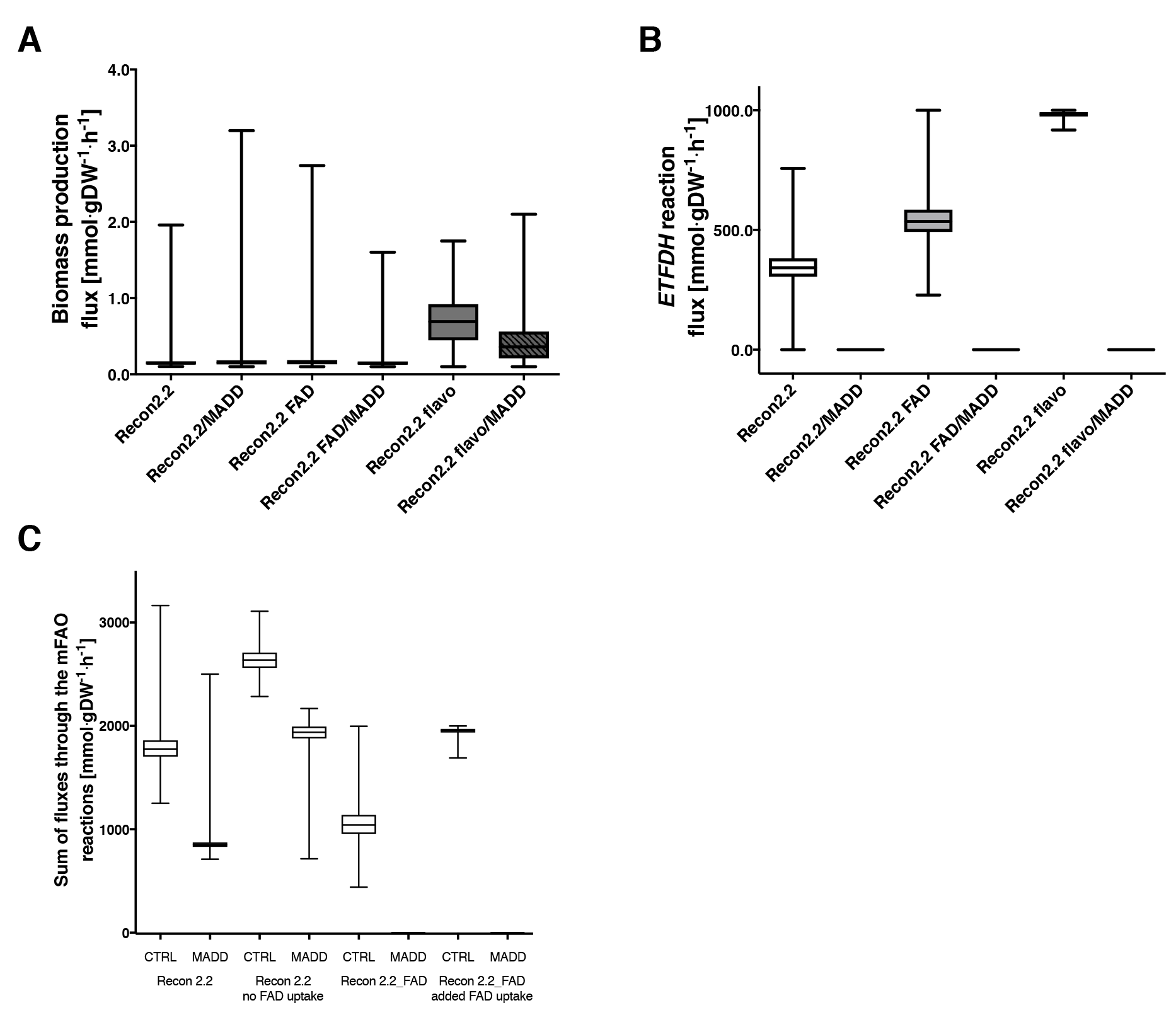
Biomass and ETFDH flux in the models used for MADD simulation. Flux values were obtained by sampling the solution space using the optGpSampler (10000 points explored, 2000 steps). Boxes span 25^th^ to 75^th^ percentiles, lines show medians, whiskers indicate minimum and maximum values. MADD was simulated by deletion of *ETFDH* in the models A. The flux via the biomass synthesis reaction remains unchanged in all models. B. None of the three models carry flux via the *ETFDH* reaction in the MADD simulation. C. Removal of FAD uptake in the Recon2.2 model does not change the incorrect model behaviour in MADD case, similarly adding a free FAD uptake to our curated model does not cause incorrect flux predictions.

**Fig S3.**
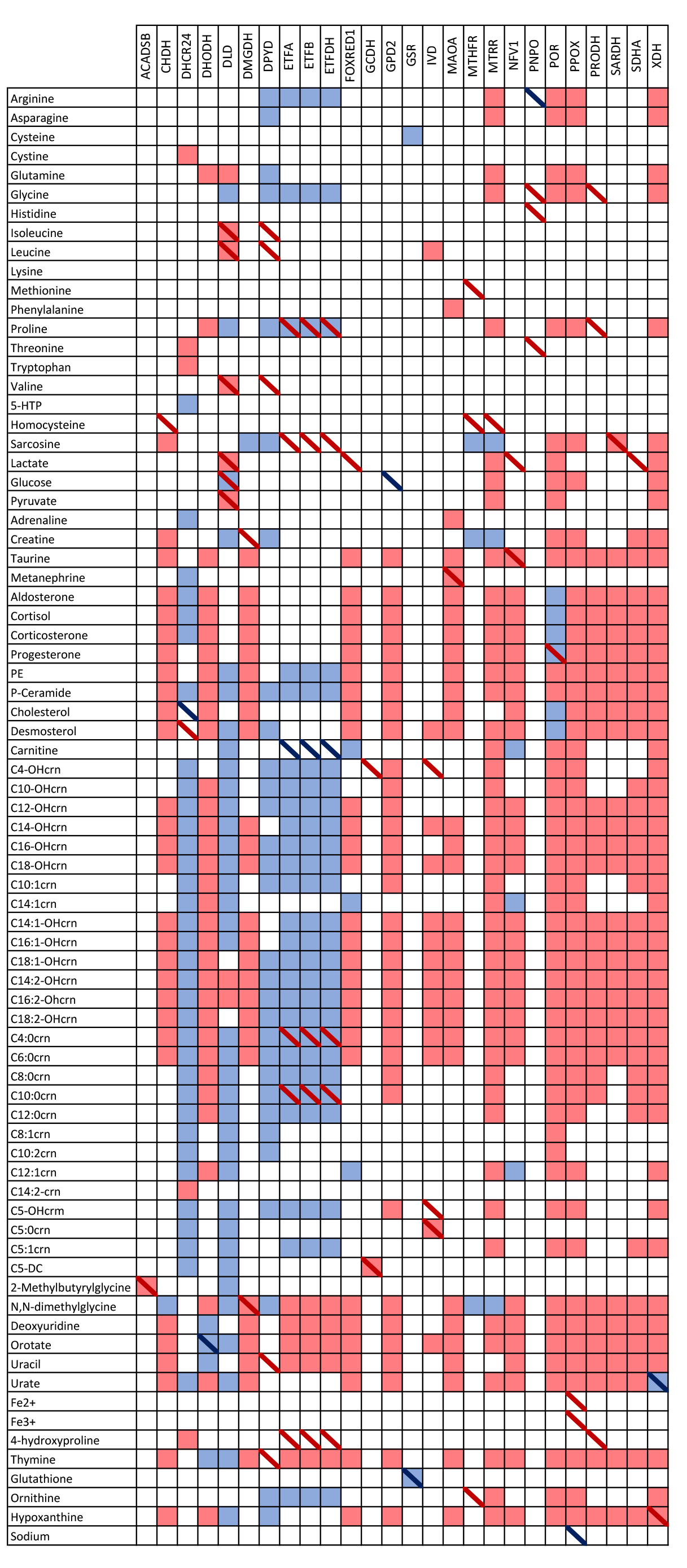
Biomarker predictions for single gene deficiencies. Detailed in Table 3 [FVA].

**Fig S4.**
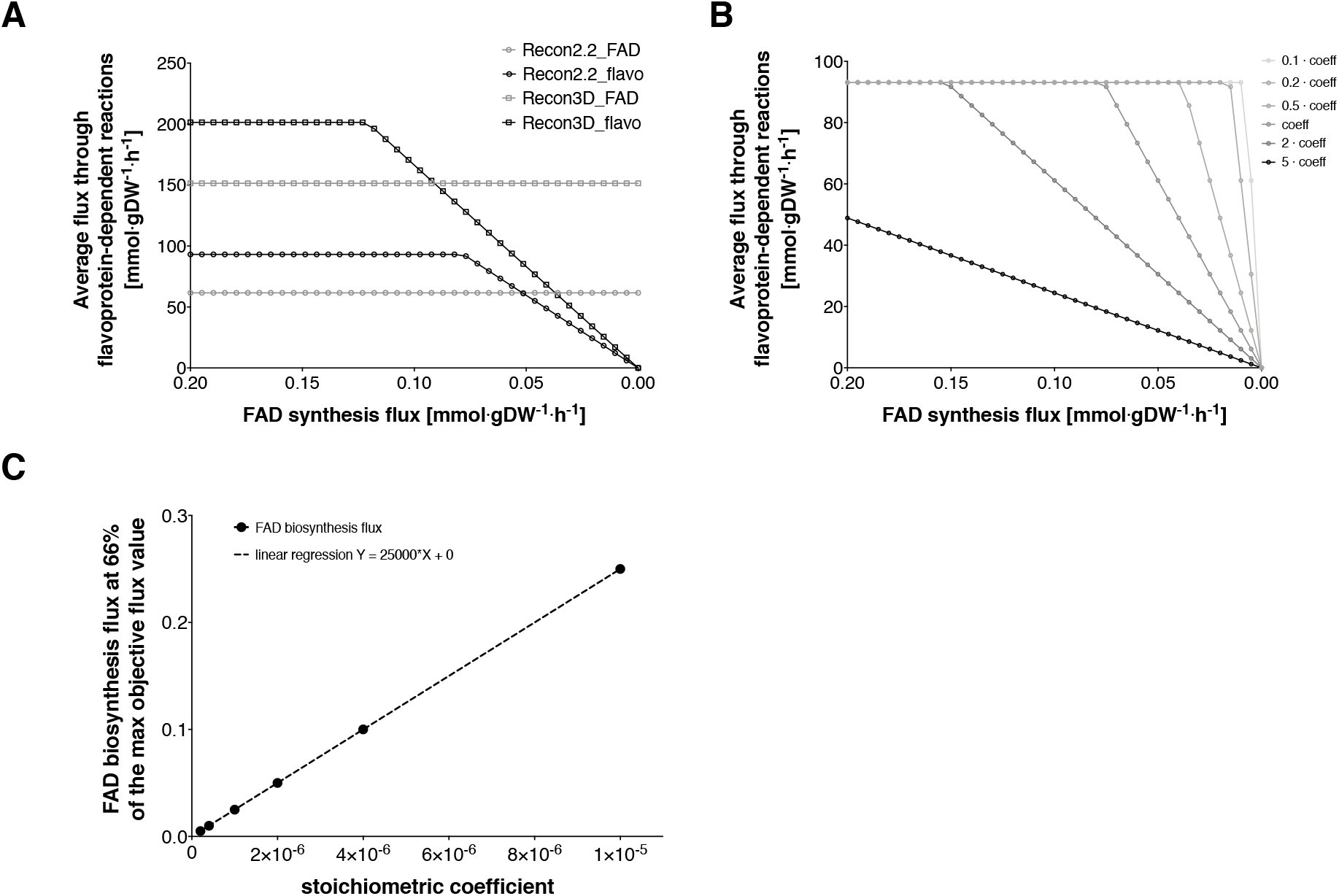
Average flux through the FAD-related reactions as a function of FAD biosynthesis flux. Maximum capacity was calculated by maximizing the objective function – a sum of fluxes through all the FAD-related reactions. A. Coupling of FAD-related reactions to FAD-biosynthesis enabled the modified Recon 3D_FAD model to respond to low cofactor availability similarly to Recon 2.2_FAD; B. Sensitivity of the average flux through flavoprotein-dependent reactions to changes in the cofactor’s stoichiometric coefficient. C. FAD biosynthesis flux required to reach the 66% of the maximal flux through the FAD-related reactions depending on the different stoichiometric coefficient of cofactor.

**Fig S5.**
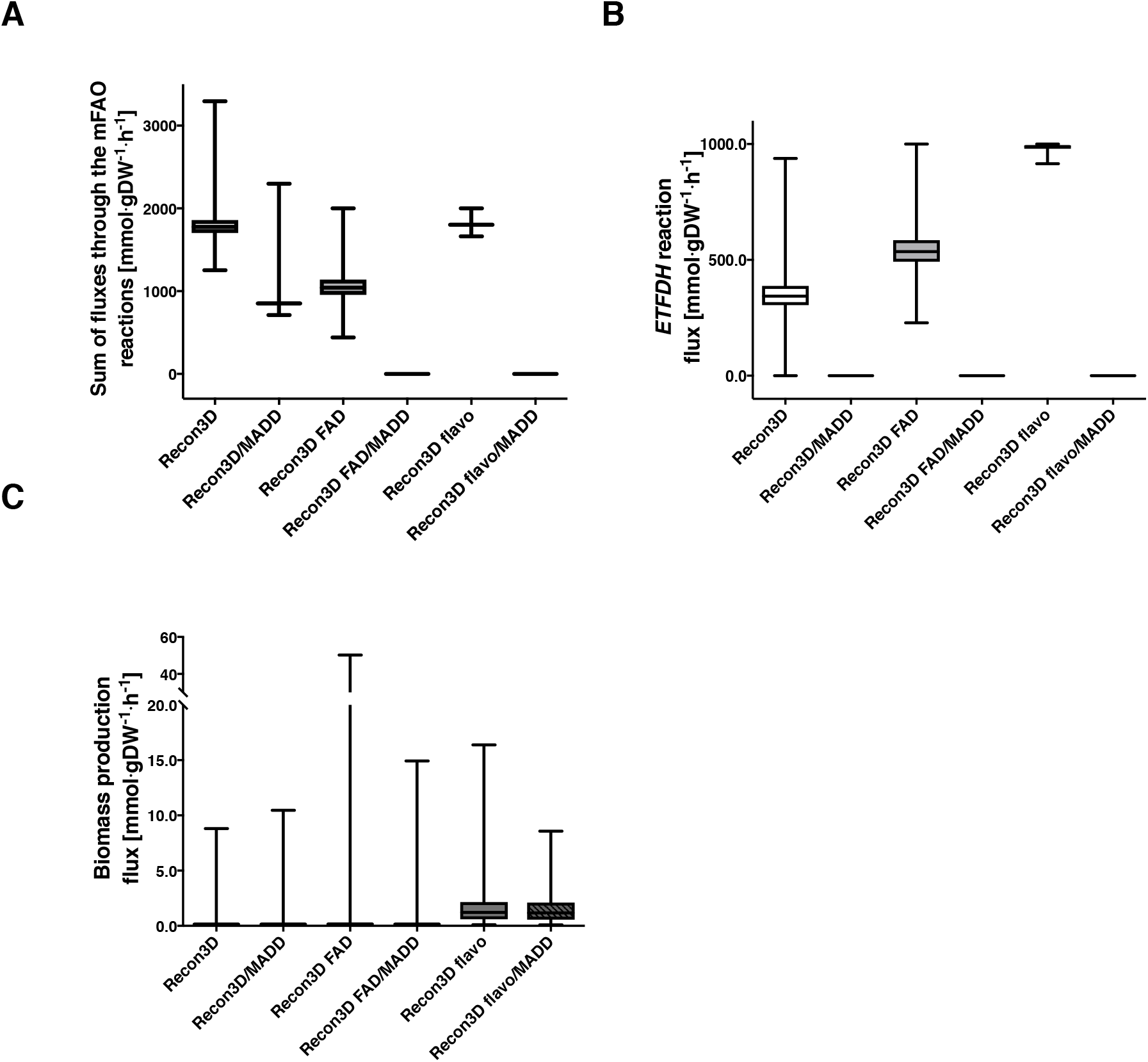
Recon 3D can correctly simulate the physiology of MADD after our FAD fix was applied. Flux values were obtained by sampling the solution space using the optGpSampler (10000 points explored, 2000 steps). Boxes span 25^th^ to 75^th^ percentiles, lines show medians, whiskers indicate minimum and maximum values. MADD was simulated by deletion of *ETFDH* in the models; A. Total flux through the mFAO (sum of fluxes through the mFAO reactions); B. None of the three models carry flux via the *ETFDH* reaction in the MADD simulation; C. The flux via the biomass synthesis reaction remains unchanged in all models.

**Table S1.** Information about human flavoproteome.

**Table S2.** Manual curation process of Recon 2.2.

**Table S3.** Metabolic biomarkers for flavoprotein-related diseases.

**Table S4.** FAD curation step applied for Recon 3D.

**Table S5.** Biomarker prediction accuracy and TPR between Recon 2.04, Recon 2.2, and Recon 3D.

**Files S1 and S2** (containing models and scripts) are available as separate files.

